# Sequencing of the MHC region defines *HLA-DQA1* as the major independent risk for anti-citrullinated protein antibodies (ACPA)-positive rheumatoid arthritis in Han population

**DOI:** 10.1101/400937

**Authors:** Jianping Guo, Tao Zhang, Hongzhi Cao, Xiaowei Li, Hao Liang, Mengru Liu, Yundong Zou, Yuanwei Zhang, Xiaolin Sun, Fanlei Hu, Yan Du, Xiaodong Mo, Xu Liu, Yue Yang, Huanjie Yang, Xinyu Wu, Xuewu Zhang, Huijue Jia, Hui Jiang, Yong Hou, Xin Liu, Yin Su, Mingrong Zhang, Huanming Yang, Jian Wang, Liangdan Sun, Liang Liu, Leonid Padyukov, Luhua Lai, Kazuhiko Yamamoto, Xuejun Zhang, Lars Klareskog, Xun Xu, Zhanguo Li

## Abstract

The strong genetic contribution of the major histocompatibility complex (MHC) to rheumatoid arthritis (RA) susceptibility has been generally attributed to *HLA-DRB1*. However, due to the high linkage disequilibrium in the MHC region, it is difficult to define the ‘real’ or/and additional independent genetic risks using the conventional HLA genotyping or chip-based microarray technology. By the capture sequencing of entire MHC region for discovery and HLA-typing for validation in 2,773 subjects of Han ancestry, we identified HLA-DQα1:160D as the strongest independent genetic risk for anti-citrullinated protein antibodies (ACPA)-positive RA in Han population (*P* = 6.16 × 10^−36^, OR=2.29). Further stepwise conditional analysis revealed that DRβ1:37N has an independent protective effect on ACPA–positive RA (*P* = 5.81 × 10^−16^, OR=0.49). The DQα1:160 coding allele *DQA1*0303* displayed high impact on joint radiographic severity, especially in patients with early disease and smoking (*P* = 3.02 × 10^−5^). Interaction analysis by comparative molecular modeling revealed that the negative charge of DQα1:160D stabilizes the dimer of dimers, leading to an increased T cell activation. The electrostatic potential surface analysis indicated that the negative charged DRβ1:37N encoding alleles could bind with epitope P9 arginine, thus may result in a decreased RA susceptibility.

In this study, we provide the first evidence that *HLA-DQA1*, instead of *HLA-DRB1*, is the strongest and independent genetic risk for ACPA-positive RA in Chinese Han population. Our study also illustrates the value of MHC deep sequencing for fine mapping disease risk variants in the MHC region.

## INTRODUCTION

Rheumatoid Arthritis (RA) is a chronic and systemic autoimmune syndrome primarily affecting peripheral joints. Results from several studies indicate that RA is a heterogeneous disease where different subsets of the disease results from complex interactions between different genetic and environmental factors^1-3^. The genetic factors are believed to influence not only disease susceptibility but also severity. The major histocompatibility complex (MHC) genes, encoding the human leukocyte antigens (HLA), represent the best described genetic risk loci linking to RA susceptibility in all populations that have been investigated so far. HLA-DR genetic variants are mainly associated with anti-citrullinated protein antibodies (ACPA)-positive RA^4-10^. Furthermore, studies have reported that the RA-risk HLA alleles are heterogeneous among ethnic groups. For example, *HLA-DRB1*04:01* is the major RA-risk alleles in European Caucasians, whereas *DRB1*04:05* is the most frequent RA-risk alleles in East Asians^8, 11-14^. In addition to *HLA-DRB1*, other HLA genes, such as *HLA-B* and *-DPB1*, have also been suggested to play a role in susceptibility to RA^9, 15^. However, due to the high linkage disequilibrium (LD) in MHC region, extended haplotype structures, and high density genes within MHC region, it is difficult to identify the ‘real’ or/and additional independent genetic risks by the conventional HLA genotyping and chip-based microarray technology, which defines MHC-resident association(s) based on indirect haplotype determination^16, 17^.

Smoking has been recognized as the most prominent environmental factor for RA development and severity^18, 19^. One important aspect of smoking as RA-risk factor is its involvement in gene-environment interaction. Smoking has greater impact on RA in individuals being carriers of *HLA-DRB1* alleles, and the interaction between smoking and *HLA-DRB1* alleles mainly confers a higher risk for ACPA-positive RA^6, 20-26^.

To fine map HLA region and identify novel variants contributing to RA, we performed a deep sequencing for entire MHC region. We analyzed HLA alleles, amino acids, SNPs, and indels across the MHC region to define the association for ACPA-positive RA. To the best of our knowledge, we showed for the first time that DQα1:160D, instead of *DRB1*0405*, is the greatest and independent risk factor for ACPA-positive RA in Han population. Conditional analysis further revealed that DRβ1:37N is an independent protective factor for ACPA–positive RA. We validated and confirmed these novel findings in an independent case-control cohort by classical HLA genotyping methodology. Moreover, we observed that one of DQα1:160D encoding alleles *DQA1*0303* confers strong risk for joint destruction in patients with early disease and smoking.

## MATERIAL AND METHODS

### Study subjects

Two independent cohorts, including 1358 subjects for discovery cohort (357 cases and 1001 controls) and 1415 subjects for validation cohort (604 cases and 811 controls), were enrolled in the study. Patients satisfied the American College of Rheumatology 1987 revised criteria for a diagnosis of RA^27^, and were recruited from the Department of Rheumatology and Immunology at Peking University People’s Hospital. All cases were ACPA-positive RA patients. ACPA were quantified using a second generation anti-CCP (anti-cyclic citrullinated peptides) antibodies ELISA kit, with a cut-off of 5 RU/mL (Euroimmun, Luebeck, Germany). Among cases, a total of 558 x-ray sets of hands were available. All x-rays were chronologically scored for assessment of bone destruction, as described previously^28, 29^.

In the discovery cohort, the healthy controls were selected by adjusting with age and sex from the control cohort of a study in psoriasis^30^. In the validation cohort, the healthy subjects were recruited from Health Care Center of People’s Hospital and were selected by adjusting with age and sex and without any disease records. All patients and healthy individuals were Han Chinese. The baseline demographic characteristics of patients and healthy controls are summarized in **Table 1** and the workflow of this study is described in **Supplementary Fig. 1**.

**Table 1.** Demographic characteristic of the study cohorts

|  | Stage I (Discovery) | Stage II (Validation) |
| --- | --- | --- |
| No. of patients/ controls | 357/1001 | 604/811 |
| **Demographic characteristics** |  |  |
| Female (%) | 80.4/ 72.4 | 81.0/82.2 |
| Age, mean ± SD years | 57.4 ± 12.1/43.0 ± 15.7 | 54.6 ± 13.0/45.3 ± 13.6 |
| **Clinical characteristics** |  |  |
| Age at onset, mean ± SD years | 47.1 ± 13.9 | 45.3 ± 14.6 |
| Disease duration, mean ± SD years | 10.3 ± 8.9 | 9.4 ± 8.7 |
| SHS, mean ± SD (n)* | 80.5 ± 63.0 (336) | 41.6 ± 51.8 (212) |

RA: rheumatoid arthritis; ACPA: anti-citrullinated proteins antibodies; SHS: modified
Sharp-van der Heijde score; SD: standard deviation.
\*: the case number when data were available.

**Figure 1.**
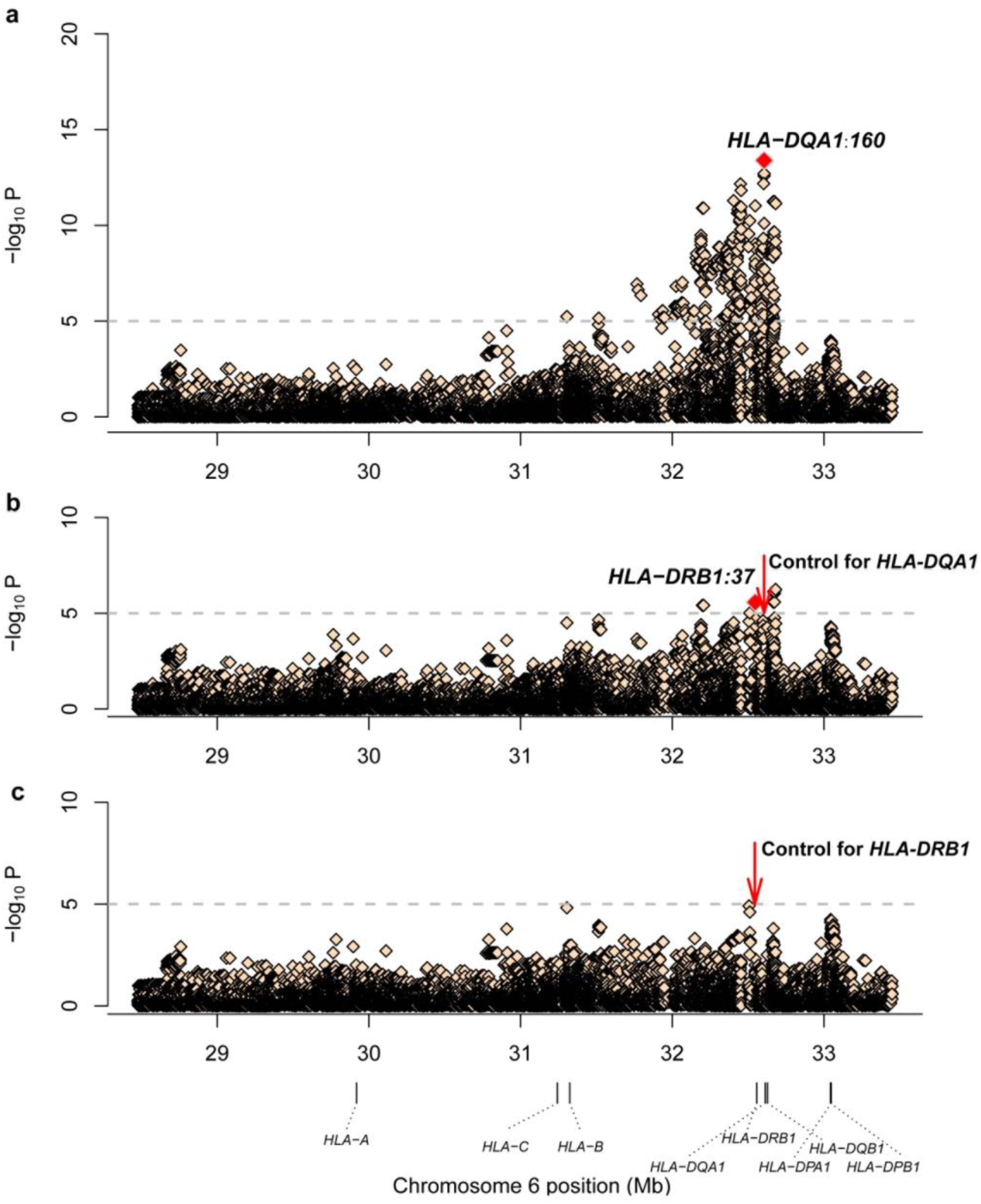
Plots of stepwise conditional analysis for ACPA–positive RA in MHC region for discovery cohort. Each diamond represents −log10(P) of the variants, including SNPs, Indels, classical HLA alleles and amino acid polymorphisms of HLA genes. The dotted horizontal line represents the suggestive significance threshold of *P* = 1 × 10^−5^. The bottom panel indicates the physical positions of the HLA genes on chromosome 6 (NCBI Build 37). (**a**) The major genetic determinants of ACPA–positive RA mapped to HLA-DQα1 corresponding Asp-160. (**b**) Subsequent conditional analyses controlling for HLA-DQα1 Asp-160 reveal an independent association at HLA-DRβ1 corresponding Asn-37. (**c**) Upon controlling for HLA-DQα1 Asp-160 and HLA-DRβ1 Asn-37, no additional significant association signal was observed.

The study was approved by the Medical Ethics Committee of Peking University People’s Hospital and informed consent was obtained from all participants.

### MHC target sequencing

In discovery stage, the MHC region was sequenced using the targeted capture sequencing methodology, as described previously^31^. Briefly, genomic DNA was extracted using DNeasy Blood & Tissue Kits (QIAGEN, 69581). Following the manufacturer’s instructions, whole genome shotgun libraries were built from 3 µg of genomic DNA (Illumina, San Diego, CA, USA). Then one µg of prepared sample library was hybridized to the capture probes for incubating at 65 °C, following the manufacturer’s protocol (Roche NimbleGen, Madison, WI, USA). The targeted fragments were subsequently captured and samples were washed twice at 47°C and at room temperature. The Platinum Pfx DNA polymerase (Roche NimbleGen, Madison, WI, USA) were used to amplify the captured fragments. The PCR products were thereafter purified and sequenced with standard 2 × 90-bp paired-end reads on the Illumina HiSeq 2000 sequencer. Sequencing data for the 1001 healthy controls were cited and selected by adjusting with age and sex from a recent publication of psoriasis project^30^.

### Alignment and variant calling

Sequenced samples were aligned to the NCBI human genome reference assembly (Build 37) using Burrows-Wheeler Aligner (BWA, version 0.5.9). On average, the MHC region was sequenced to a mean depth of 94 X, with 96.6% covered by at least one read and 93.1% covered by at least ten reads for cases. The data summary of the health controls is described in the previous study^30^. Then using SAMtools (v0.1.17), the file was conversed from SAM to BAM, the sorted and indexed BAM files were generated and duplicates were marked. To perform the realignment around known Indels, the BAM files were analyzed using the Genome Analysis Toolkit (GATK v1.4). All aligned read data were subjected to CountCovariates (GATK) on the basis of known single-nucleotide variants (SNVs) (dbSNP135) and TableRecalibration (in GATK) was used to recalibrate the base quality. Single nucleotide variants and indels were called jointly with GATK UnifiedGenotyper. Then, the GATK resource bundle was used for variant quality score recalibration, which includes known SNP sites from HapMap v3.3, dbSNP135, the Illumina Omni2.5 array, the Mills and the 1000G gold standard Indels as training data ^32^. To build a genotype matrix as input for the subsequent analysis, the genotypes for each detected variant position were extracted from all samples.

### HLA typing

In discovery cohort, a total of 26 highly polymorphic HLA genes were genotyped according to the Short Oligonucleotide Analysis Package (SOAP)-HLA^31^. SOAP-HLA is a flow of sequencing data analysis pipeline to type any HLA genes using capture sequenced data based on IMGT/HLA database with a high accuracy^31^. The following HLA genes were typed including *HLA-A, HLA-B, HLA-C, HLA-E, HLA-F, HLA-G, HLA-H, HLA-J, HLA-K, HLA-L, HLA-P, HLA-V, HLA-DRA, HLA-DRB1, HLA-DQA1, HLA-DQB1, HLA-DPA1, HLA-DPB1, HLA-DMA, HLA-DMB, HLA-DOA, HLA-DOB, HLA-MICA, HLA-MICB, HLA-TAP1*, and *HLA-TAP2*. The amino acid sequence of each HLA allele was determined according to the IMGT/HLA database (Release 3.22.0).

In validation cohort, a total of 1415 individuals (604 cases and 811 controls) were genotyped for *HLA-A, -B, DRB1, -DPB1* and -*DQA1*. Genotypes of *HLA-A, -B, -DRB1* and *-DPB1* alleles were determined by using the next generation sequencing (NGS) method^33^. In brief, the amplicons were pooled and sheared randomly. Gel slicing was used to recover the sequencing libraries with fragments which include the library adapters between 400 and 700 bp in length. Then, the sequencing was performed on Illumina Miseq with PE 150 bp reads in a single run. Finally, using the unique variant calling and haploid sequencing assembly algorithm with the short sequence read as input, the genotypes were accurately obtained. *HLA-DQA1* were genotyped by sequencing of both exons 2 and 3 using the gold standard Sanger sequencing method^34, 35^.

### Quality control

Sequencing data were evaluated against a quality control metrics for all the samples. We restricted each individual as follows: (i) average sequencing depth ≥4X; (ii) 90% of the target region covered by 4X; (iii) GC content within 42%-48%. According to the criteria, a total of 58 samples from the discovery stage were filtered and removed for further analysis (**Supplementary Table 1**).

After initial sample quality control for the MHC capture sequencing, we performed further filtering to identify the high-confidence SNPs and Indels in targeted region. Following criteria were applied: i) pass ratio ≥ 0.9 (Q100 and Q500 were defined as pass for SNPs and Indels, respectively); ii) missing rate ≤ 0.1 (a depth of ≥ 5X was considered as high quality; individuals failed to meet the criteria were considered as missing); iii) minor allele frequency (MAF) ≥ 0.01; iv) Hardy-Weinberg test *P-* value ≥ 1.0 × 10^−6^ (**Supplementary Table 2**). For HLA types and amino acids, the same process was performed except for the pass ratio criteria (**Supplementary Table 2**).

### Statistical analyses

Using the same data processing procedure and analysis^30, 36^, logistic regression model was applied to test the association between ACPA-positive RA and the variants in the MHC region, adjusting by gender. We define the HLA variants by including the four digit biallelic classical HLA alleles, biallelic SNPs, Indels, and biallelic HLA amino acid polymorphisms for respective residues within MHC region. The top five principal-components (PCs) were applied to control for population stratification in the discovery study. For the individual HLA allele and amino acid variant, the association was determined after stratifying the data using the relative predispositional effect (RPE) method^37^. Thus, the logistic regression model is as follows:

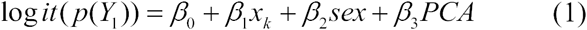

Where Y_1_ is RA status (1 if ACPA-positive RA and 0 otherwise) and *x*_*k*_ is the genotype at the *k*th variants. The *β*_*0*_ is the logistic regression intercept and *β*_*1*_, *β*_*2*_, *β*_*3*_ are the effect size of *k*th variants, gender and the PCA respectively. The P values for this test were observed while the odds ratio (OR) and standard error (SE) of OR were estimated.

To assess the independent effects of candidates identified in logistic analysis, we assumed the logistic regression model additionally including the most significant loci as covariates for the stepwise conditional regression analysis. If additional independent risk factors were identified, we further consecutively included them as covariates in the subsequent multivariate analyses in a forward conditional stepwise manner until none of loci met the cut off *P*-value^30, 38^. Assuming that there may be linkage disequilibrium (LD) between the intergenic regulatory variants and specific genes, thus the independence of intergenic variants from its surrounding genes should be tested^36^. If the *P*-value of the intergenic variants is no longer less than the significant threshold after conditioning on the polymorphism sites (including SNPs, INDELs, amino acids, HLA type) of the nearby genes, we should perform condition analysis on its tagged genes, otherwise the intergenic variant is considered a real independent association locus. The unpaired T-test was applied to assess the significance of differences in radiological scores between two groups.

In discovery stage, a *P*-value less than 1.0 × 10^−5^ was suggested as significance threshold. In validation and combined stages, a conventional genome-wide statistical significance threshold of *P* less than 5.0 × 10^−8^ was applied. Analysis of radiological severity was conducted in R statistics program.

### Comparative modeling

The comparative modeling was conducted using Modeller v9.14^39^. The crystal structure of HLA-DR1 (*DRA-DRB1*0101*) (PDB ID: 1AQD) was employed as template for comparative modeling of *DQA1*0303-DQB1*0401*^40^. *The overall sequence identify and similarity between DQA1*0303-DQB1*0401* and HLA-DR1 are 61.7% and 76.6%, respectively. The crystal structure of HLA-DR3 (*DRA-DRB1*03:01*) (PDB ID: 1A6A) has been solved and thereby was used as template for the comparative modeling of *DRA-DRB1*13:0*1 and *DRA-DRB1*13:02*^41^. The sequence identities of HLA-DR3 to *DRA-DRB1*13:01* and *DRA1-DRB*13:02* were 98.1%, and 97.8%, respectively. For each comparative modeling, ten models were generated, and the structure with the lowest probability density function total energy was selected for structural refinement. Energy minimization was performed using the Amber14 package with the Amber ff14SB force field^42^. Each structure model was solvated in an octahedron TIP3P water box and neutralized by adding proper counter ions. The distance of box boundary and structure model was set to 10 Å. The particle-mesh-Ewald (PME) method^43^ was used for the treatment of long-range electrostatic interactions. The non-bond interaction cutoff was set to 8.0 Å. Each simulation system was subjected to three stages of energy minimization, including (1) 5000 steps of steepest descent (SD) and 2000 steps of conjugate gradient (CG) minimization with harmonic restraints (10 kcal/Å) applied on all structural atoms; (2) 5000 steps of SD minimization and 2000 steps of CG minimization with reduced harmonic restraints (2 kcal/Å) on backbone atoms; (3) 10,000 steps of steepest descent and 5000 steps of conjugate gradient minimization with all restraints removed.

## RESULTS

### HLA-DQα1:160D is the strongest genetic risk for ACPA–positive RA – Discovery by MHC sequencing

To define the independent association(s) and/or discover any novel variant(s) contributing to RA in addition to *HLA-DRB1*, we first conducted a capture sequencing of the whole MHC region in 357 patients with ACPA–positive RA and 1001 previously sequenced healthy controls of Han Chinese. After quality control, a total of 24,177 variants, 166 HLA types, and 1,283 amino acids were obtained (**Supplementary Table 2**).

In total, 563 variants (including HLA types and amino acids) showed significant associations by a cut-off of *P*-value less than 1.0 × 10^−5^(**Supplementary Table 3**). We found that the top association signal was mapped to *HLA-DQA1*, with a peak at HLA-DQα1 amino acid position 160 (DQα1:160D, *P* = 4.03 × 10^−14^, OR=2.42, 95% CI 1.92-3.04), followed by DQα1:160A (*P* = 2.01 × 10^−13^, OR=2.34, 95% CI 1.86-2.93) and *DQA1*0303* (*P* = 2.62 × 10^−13^, OR=3.02, 95% CI 2.25-4.06) (**Fig. 1** and **Supplementary Table 3**). *HLA-DRB1*0405* allele, the putative greatest RA risk in Asians in previous reports, also showed a strong association with ACPA-positive RA, but fell out of the top 10 risk variants (*P* = 9.48 × 10^−12^, OR=3.22, 95% CI 2.30-4.50) (**Supplementary Table 3**). These results indicated that by sequencing the entire MHC region we discovered HLA-DQα1:160D may be the top genetic risk for ACPA–positive RA in Han population, instead of the well-known *HLA-DRB1 *0405*.

### HLA-DRβ1:37N has an independent protective effect on ACPA–positive RA

By stepwise conditional analysis on HLA-DQα1:160D, the second signal was mapped to a list of SNPs, with a peak at rs7764856 (chr6_32680640_T_A) (**Supplementary Table 4**). The SNPs are in high LD (r^2^= 0.72) and all located in intergenic region between *DQA2* and *DQB1* according to NCBI RefSeq database (http://www.ncbi.nlm.nih.gov/RefSeq). An immediate significant association was observed for HLA-DRβ1:37N after these intergenic SNPs (**Fig. 1** and **Supplementary Table 4**). Of note, HLA-DRβ1:37N has an independent protective effect on ACPA–positive RA (*P* = 2.71 × 10^−6^, OR=0.51, 95% CI 0.39-0.68). Conditioning on DRβ1:37N, the intergenic SNPs lost their association and no additional independent association(s) reached the suggestive statistical significance threshold of *P* < 1.0 x10^−5^ (data not shown).

**Table 2.**
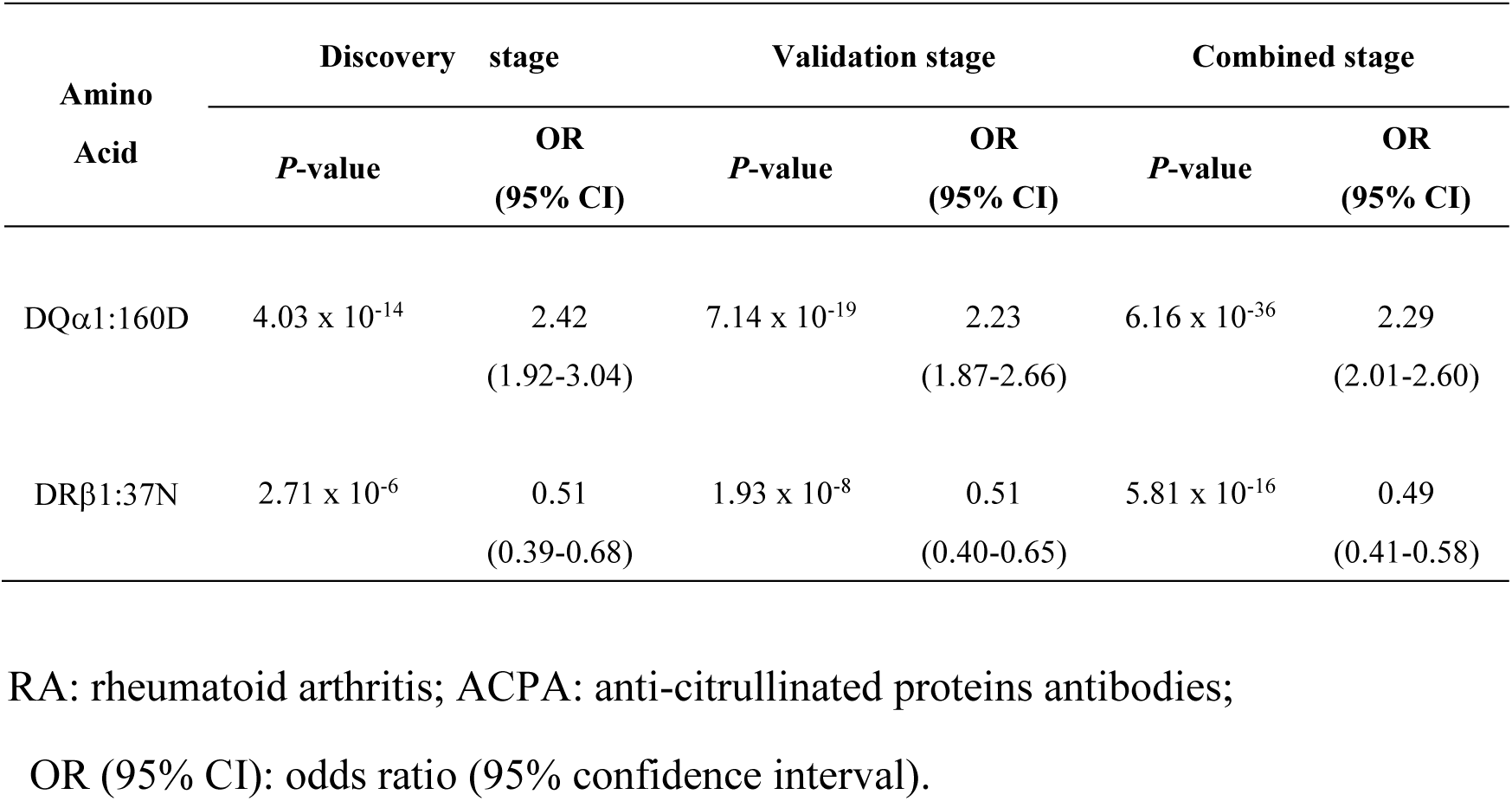
Independent effects of DQα1:160D and DRβ1:37N in ACPA-positive RA

To assess whether DQα1:160D and DRβ1:37N were independent of each other, we performed the conditional analysis starting from DRβ1:37N. As shown in **Supplementary Fig. 2**, after conditioning on DRβ1:37N, DQα1:160D displayed even stronger association (*P* = 1.19 × 10^−18^, OR=3.53, 95% CI 2.67-4.68). This indicates that DQα1:160D also has an independent effect on ACPA–positive RA.

### The validation study confirms the findings discovered by capture sequencing

To validate the findings discovered by capture sequencing, we performed Sanger or NGS to genotype *HLA-A, -B*, -*DRB1*, -*DQA1*, and -*DPB1* genes in an independent case-control cohort, consisting of 604 cases with ACPA–positive RA and 811 healthy subjects. Joint analysis was then performed by combining the results from discovery and validation cohorts.

In line with the findings in the discovery stage, DQα1:160D showed consistent top association with ACPA–positive RA in the validation panel (*P* = 7.14 × 10^−19^, OR = 2.23, 95% CI 1.87-2.66), followed by *DQA1*0303* (*P* = 5.74 × 10^−18^, OR = 3.13, 95% CI 2.41-4.05) and DQα1:160A (*P* = 4.18 × 10^−17^, OR=2.10, 95% CI 1.77-2.50). A consistent association was also observed for *DRB1*0405* (*P* = 5.76 × 10^−15^, OR = 3.40, 95% CI 2.50-4.62) (**Supplementary Table 5**). After conditioning to DQα1:160D, though DRβ1:96H became the second independent signal (*P* = 2.80 × 10^−10^, OR = 1.68, 95% CI 1.43-1.97), it was followed by DRβ1:37N (*P* = 1.93 × 10^−8^, OR=0.51, 95% CI 0.40-0.65) (**Supplementary Table 6**). When conditioning on DRβ1:37N, the DRβ1:96H lost the association according to the genome-wide statistical significance threshold of *P* < 5.0 × 10^−8^ (*P* = 1.18 × 10^−4^, OR = 1.45, 95% CI 1.20-1.76, data not shown).

Joint analysis of discovery and replication panels provided compelling evidence that HLA-DQα1:160D conferred the highest risk on ACPA–positive RA (*P* = 6.16 × 10^−36^, OR=2.29, 95% CI 2.01-2.60), followed by DQα1:160A (*P* = 3.29 × 10^−33^, OR=2.17, 95% CI 1.91-2.47) and *DQA1*03:03* (*P* = 5.13 × 10^−33^, OR=3.17, 95% CI 2.63-3.83) (**Fig. 2**, **Supplementary Table 7**, and **Table 2**). After conditioning to DQα1:160D, though the DRβ1:96H remained as the second independent signal (*P* = 4.90 × 10^−16^, OR=1.64, 95% CI 1.45-1.84, **Fig. 2 and Supplementary Table 7**), DRβ1:37N also displayed strong association (*P* = 5.81 × 10^−16^, OR=0.49, 95% CI 0.41-0.58, **Fig. 2**, **Supplementary Table 7**, and **Table 2**). When conditioning on DRβ1:37N, there was no additional independent association(s) reached the study-wide statistical significance of *P* < 5.0 × 10^−8^ (data not shown).

**Figure 2.**
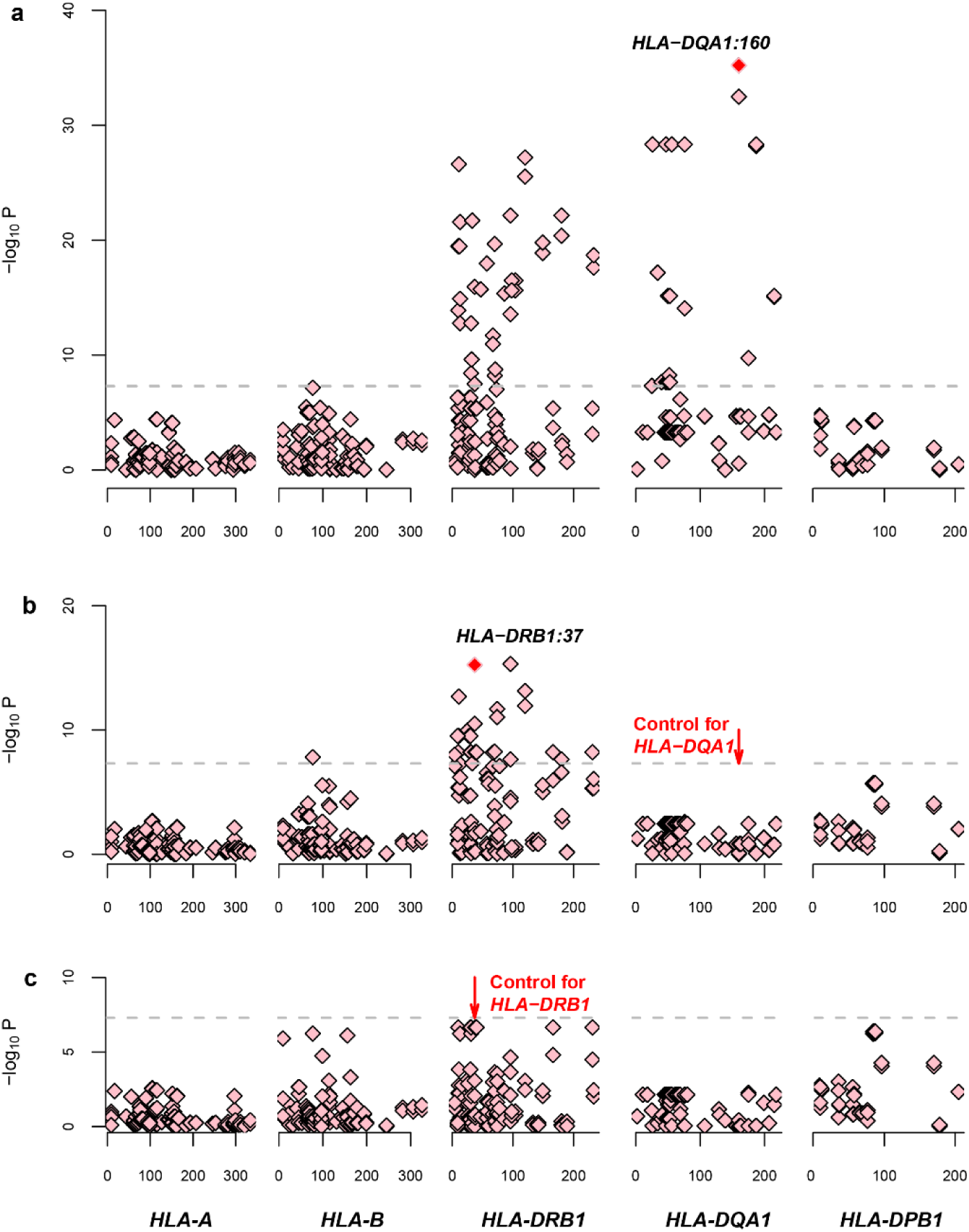
Joint analysis of discovery and replication panels in the MHC region. The association for each locus used for conditioning (*HLA-DQA1, HLA-DRB1*) is shown in red in each panel. For each panel, the horizontal axis shows the position of amino acid for each HLA gene and the vertical axis shows negative log10-transformed *P* values for association. The dashed horizontal line corresponds to the significance threshold of *P*=5×10^−8^.

### Exclusive dissection of *HLA-DRB1* indicates that DRβ1 variants could be strong risks for ACPA-positive RA, if the effect of *HLA-DQA1* is ignored

As mentioned above, *HLA-DRB1* was recognized as the strongest RA risk in previous studies, especially for ACPA-positive RA and variations *DRB1*0405*, DRβ1:11, 13, 57, 74, and 71 have been shown to confer risks for RA in Asian patients ^10^. Thus, we next investigated RA association at *HLA-DRB1* separately. As shown in **Supplementary Table 8**, multiple alleles at *HLA-DRB1* showed strong associations, with DRβ1:120N (*P* = 6.46 × 10^−28^, OR=2.27, 95% CI 1.96-2.63), *DRB1*0405*(*P* = 6.55 × 10^−28^, OR=3.40, 95% CI 2.73-4.23), and DRβ1:11V (*P* = 2.43 × 10^−27^, =2.45, 95% CI 1.94-2.60) being the top three risks. When conditioning on either DRβ1:11V or :120N, the second signal was seen for DRβ1:31I (*P* = 2.23 × 10^−18^, OR=1.93, 95% CI 1.67-2.24) and DRβ1:13F (*P* = 2.90 × 10^−17^, OR=1.82, 95% CI 1.58-2.09) (**Fig. 3**). After further conditioning to either DRβ1:13F or :31I, strong association signals were observed for *DRB1*04:05* (*P* = 6.26 × 10^−11^, OR=2.53, 95% CI 1.92-3.35) and DRβ1:57S (*P* = 3.14 × 10^−9^, OR=1.81, 95% CI 1.49-2.20), respectively. Then we continued conditioning on DRβ1:57S, DRβ1:74A showed a suggestive association with ACPA-positive RA (*P* = 5.09 × 10^−7^, OR = 0.72, 95% CI 0.63-0.82). When conditioning on DRβ1:74A, DRβ1:71E also showed a suggestive association (*P* = 2.95 × 10^−6^, OR=0.37, 95% CI 0.24-0.56). Our results indicated that if the effect of *DQA1* is ignored, *DRB1*0405*, amino acid variants at DRβ1:11, 13, 57, 74, and 71 can come up and be very strong risk factors for ACPA-positive RA.

Next, we compared the individual amino acid frequencies between Han Chinese and European populations. As shown in **Fig. 4A**, both DQα1:160D and DQα1:160A are common amino acids in Han Chinese and were significantly increased in ACPA-positive RA patients. However, the two amino acids have not been detected in European population^9^. For amino acid positions 11, 13, 57, 71 and 74 in DRβ1, the amino acid frequencies are similar between two ethnic groups or case and controls, except for a few amino acids. For example, DRβ1:11D, 13G, 57V, and 74E are common in Han Chinese but rare in Europeans (**Fig. 4B, C, D, F**). In contrast, DRβ1:71K is rare in Han Chinese but common in Europeans (**Fig. 4E**).

**Figure 3.**
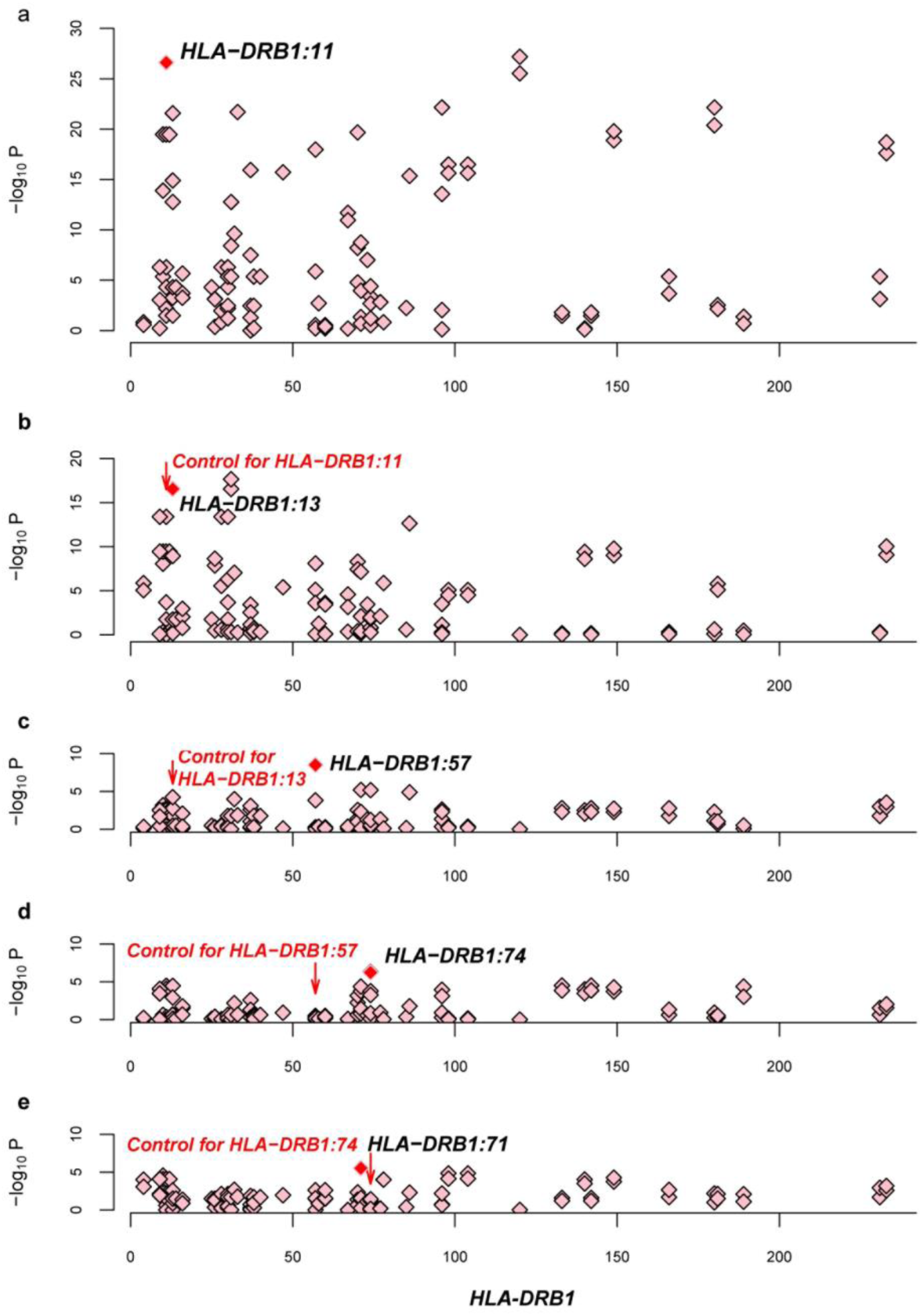
Plots of stepwise conditional analysis for HLA-DRβ1 in combined cohort. (**a**) Amino acid position 120 represents the strongest association with ACPA– positive RA (*P* < 10^−27^), followed by position 11 (*P* < 10^−26^). (**b**) Controlling for position 11 or 120, position 13 is an independent risk for ACPA–positive RA (*P* = 2.90 × 10^−17^). (**c**) Controlling for positions 11 and 13, position 57 showed an independent association (*P* = 3.14 × 10^−9^). (**d**) Controlling for positions 11,13 and 57, position 74 becomes a suggestive signal (*P* = 5.09 ×10^−7^) (**e**) Conditioning on positions 11,13, 57 and 74 revealed a suggestive association for amino acid 71 (p=2.95 ×10^−6^).

**Figure 4.**
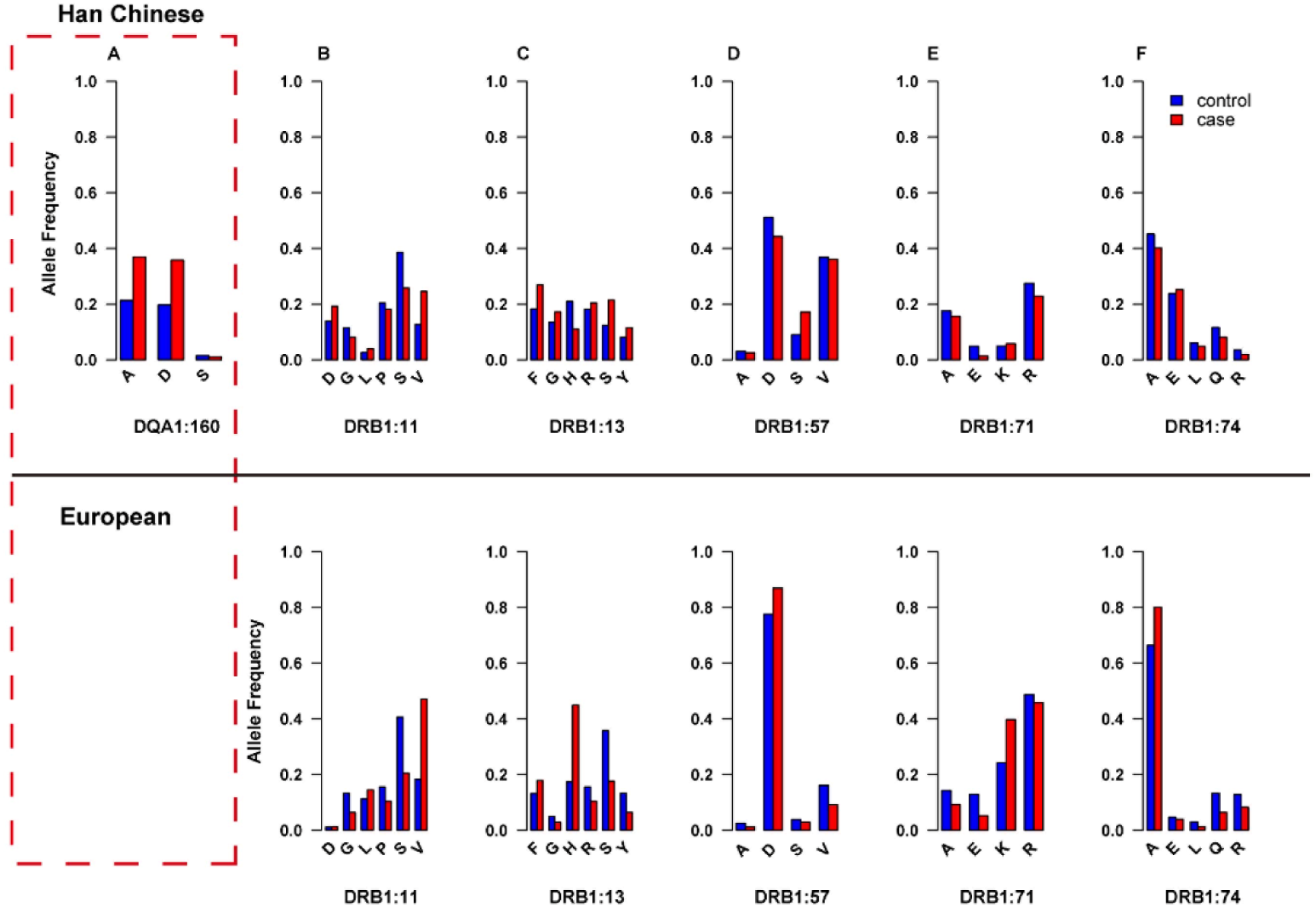
Comparison of individual amino acid frequencies within DQα1:160 and DRβ1:11, 13, 57, 71, and 74 in Han Chinese and European populations. The individual amino acid frequencies are plotted in healthy controls (blue) and cases (red). Upper panel shows the amino acid frequencies in Han Chinese population (the data derived from present study). Lower panel shows the amino acid frequencies in European population (the data cited from Raychaudhur’s study^9^).

### *DQA1*0303*, an allele encoding DQα1:160D, confers increased risk of joint damage in early ACPA-positive RA

DQα1:160D is encoded by two alleles, i.e. *DQA1*0302* and *\*0303*. We next examined whether the top susceptible factor DQα1:160D and its encoding alleles confer a risk for the severity of joint damage in ACPA-positive RA. A total of 557 patients with available SHS data were divided into three groups according to the disease durations (≤ 1 years, 1–10 years, or ≥ 10 years). Overall, there was no difference in SHS according to either DQα1:160 variations or its coding allele polymorphisms in three disease stages (data not shown). As smoking is a well-established environmental factor contributing to ACPA-positive RA susceptibility and severity, we further stratified the patients by smoking status. Though there was no significant difference in SHS between DQα1:160D carriers and non-carriers after stratifying by smoking status, in the early disease stage one of its coding allele *DQA1*0303* showed high impact on radiographic scores in smoking group (*P* = 3.02 × 10^−5^). Similarly, in early disease stage *DQA1*0303* carriers with smoking had higher radiographic scores than *DQA1*0303* carriers without smoking (*P* = 4.05 × 10^−8^, **Fig. 5a**). In the early disease stage *DRB1*0405* also showed a higher impact on radiographic score in smoking group (*P* = 3.02 × 10^−5^). *DRB1*0405* carriers with smoking had increased radiographic scores, compared to *DRB1*0405* carriers without smoking (*P* = 6.96 × 10^−6^, **Fig. 5b**). These findings are consistent with previous finding that the gene–environment interaction between *DRB1* variants and smoking contribute to ACPA–positive RA. Our data also suggest that DQα1:160 coding allele *DQA1*0303* has high impact on radiographic severity of ACPA-positive RA, especially in patients with early disease and smoking.

**Figure 5.**
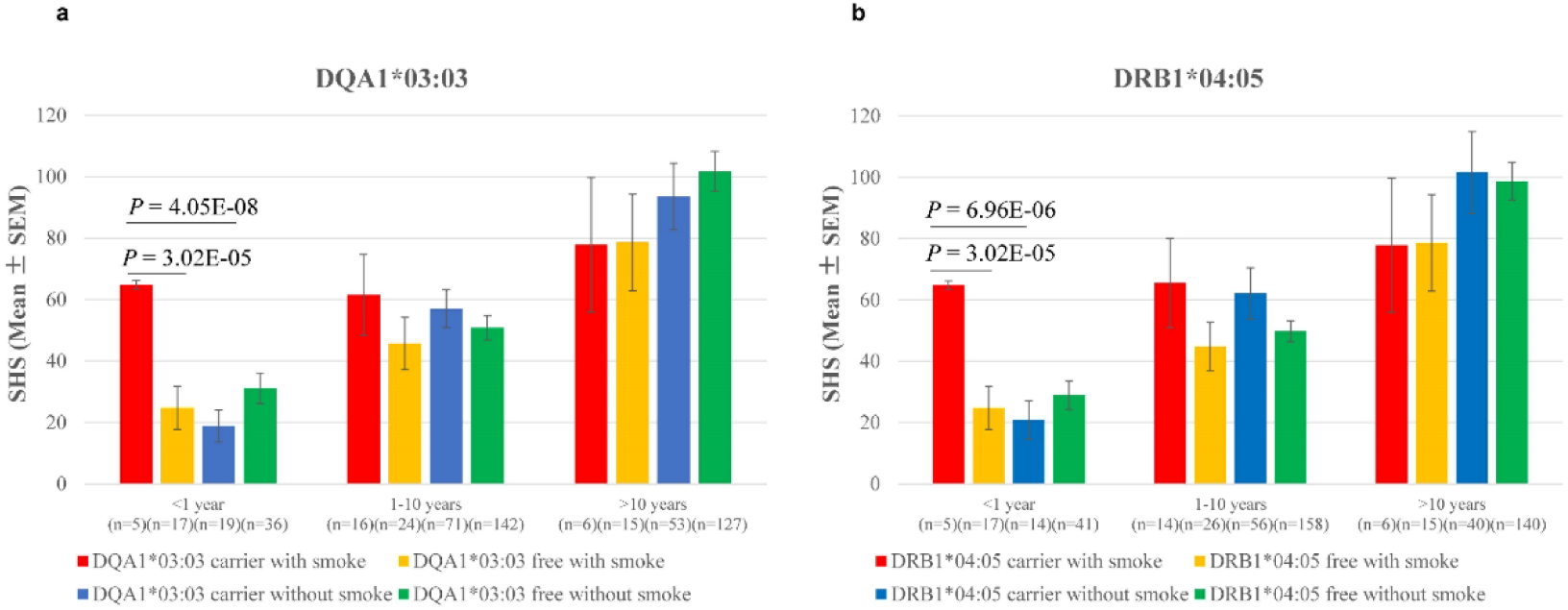
Impact of *DQA1*0303* on risk of joint damage in ACPA-positive RA. (a) In early disease stage *DQA1*0303* showed high impact on radiographic scores in smoking group (*P* = 3.02 × 10^−5^). Similarly, *DQA1*0303* carriers with smoking had higher radiographic scores than DQA1*0303 carriers without smoking (*P* = 4.05 × 10^−8^). (b) In early disease stage *DRB1*0405* also showed a high impact on radiographic score in smoking group (*P* = 3.02 × 10^−5^). *DRB1*0405* carriers with smoking had increased radiographic scores, compared to *DRB1*0405* carriers without smoking (*P* = 6.96 × 10^−6^).

### Additional negative charge of D160α enhances the interaction with DQβ1, leading to an increased T cell activation

MHC class II molecules could present in the form of either heterodimers or dimer of dimers^40, 44, 45^. The dimer of dimers appears to play an important role in T cell response to low affinity antigens by enhancing overall affinity between MHC/peptide and TCR^45, 46^. The electrostatic interactions between the interface residues are critical for maintaining structural stability as well as T cell activation^47^. According to sequence analysis, D160α locates far away from antigen binding groove and should impose little influence on epitope binding. To investigate the potential function of DQα1:160D, we constructed the dimer of dimer structure for *DQA1*0303-DQB1*0401* by comparative modeling, as it is the major D160α encoding haplotype in Han Chinese (69.0% from our data and 56% in the Han MHC database^30^). As shown in **Fig. 6A**, D160α is adjacent to the dimer of dimer interface and may contribute to the stability of dimer of dimers. Strong electrostatic interactions are observed between negative DQα1 interface residues (D161α, E182α and E184α) and positive DQβ1’ interface residues (R105β, H111β and H112β). Compared with non-charged A160α or S160α coded by other *DQA1* alleles, the additional negative charge introduced by D160α further enhances the interaction with DQβ1 in the other dimer, which may lead to an increased T cell activation (**Fig. 6B**).

**Figure 6.**
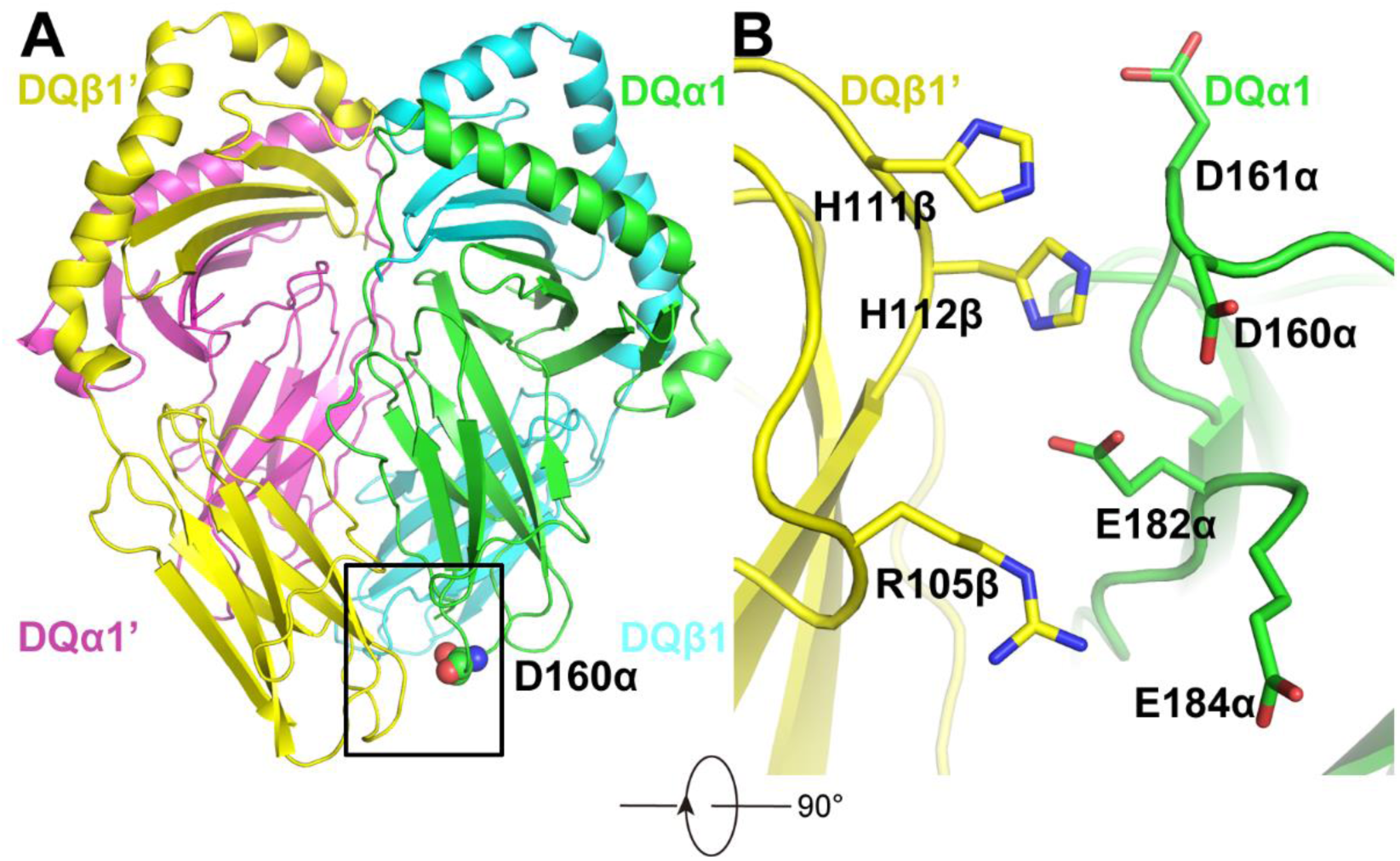
The dimer of dimer structure of *DQA1*0303-DQB1*0401*. (A) Overall dimer of dimer structure of *DQA1*\*0303-*DQB1*\*0401. One dimer is composed of DQα1 and DQβ1, whereas the other dimer is composed of DQα1’ and DQβ1’. The DQα1, DQβ1, DQα1’ and DQβ1’ were shown as green, cyan, purple, and yellow cartoon, respectively. (B) The interface of dimer of dimer structure. DQα1 and DQβ1’ residues involved in the interaction are displayed sticks and their carbon atoms are colored in green and yellow, respectively, whereas nitrogen and oxygen atoms are colored in blue and red, respectively.

### The negatively charged P9 pockets from DRβ1: 37N encoding alleles benefit electrostatic interaction and epitope P9 arginine binding

It is well accepted that *HLA-DRB1* susceptibility alleles are strongly associated with ACPA-positive RA, and its encoded molecules preferentially present the citrullinated autoantigens^48^. The susceptibility is determined by the electrostatic property of antigen binding pockets and the ability to differentially recognize citrullinated antigens. The positively charged antigen binding pocket in RA-susceptible alleles preferentially accommodate citrullinated antigens, whereas electronegative or electroneutral pockets in RA-resistant alleles can bind both arginine and citrullined antigens^49^. The amino acid residues at position 37 in DRβ1 are located within the P9 pocket of antigen binding groove. By sequence analysis, we found four alleles containing an asparagine at position DRβ1:37 (Asn37β or 37N), including *DRB1*03:01, *09:01, *13:01*, and *\*13:02*. Though *DRB1*0901* was reported to be the risk allele for RA in Koreans^11^, its influence on developing of ACPA-positive RA subgroup seemed protective^50, 51^. We then constructed the structure models for other allele-containing haplotypes by comparative modeling. The electrostatic potential surface and the residues of the P9 pocket from the three models are shown in **Fig. 7**, respectively. *DRA1-DRB1*03:01, DRA1-DRB1*13:01* and *DRA1-DRB1*13:02* share identical P9 pocket, which is composed of Asn69α, Met73α, Tyr30β, Asn37β, Val38β and Asp57β (**Fig. 7D, E** and **F**). The electrostatic potential surface analysis indicates that P9 pockets of *DRA1-DRB1*03:01, DRA1-DRB1*13:01* and *DRA1-DRB1*13:02* are negatively charged and could benefit electrostatic interaction with epitope P9 arginine (**Fig. 7A, B** and **C**). Collectively, all three alleles containing negatively charged P9 pocket of Asn37 could bind the epitope P9 arginine and thus may result in a decreased RA susceptibility.

**Figure 7.**
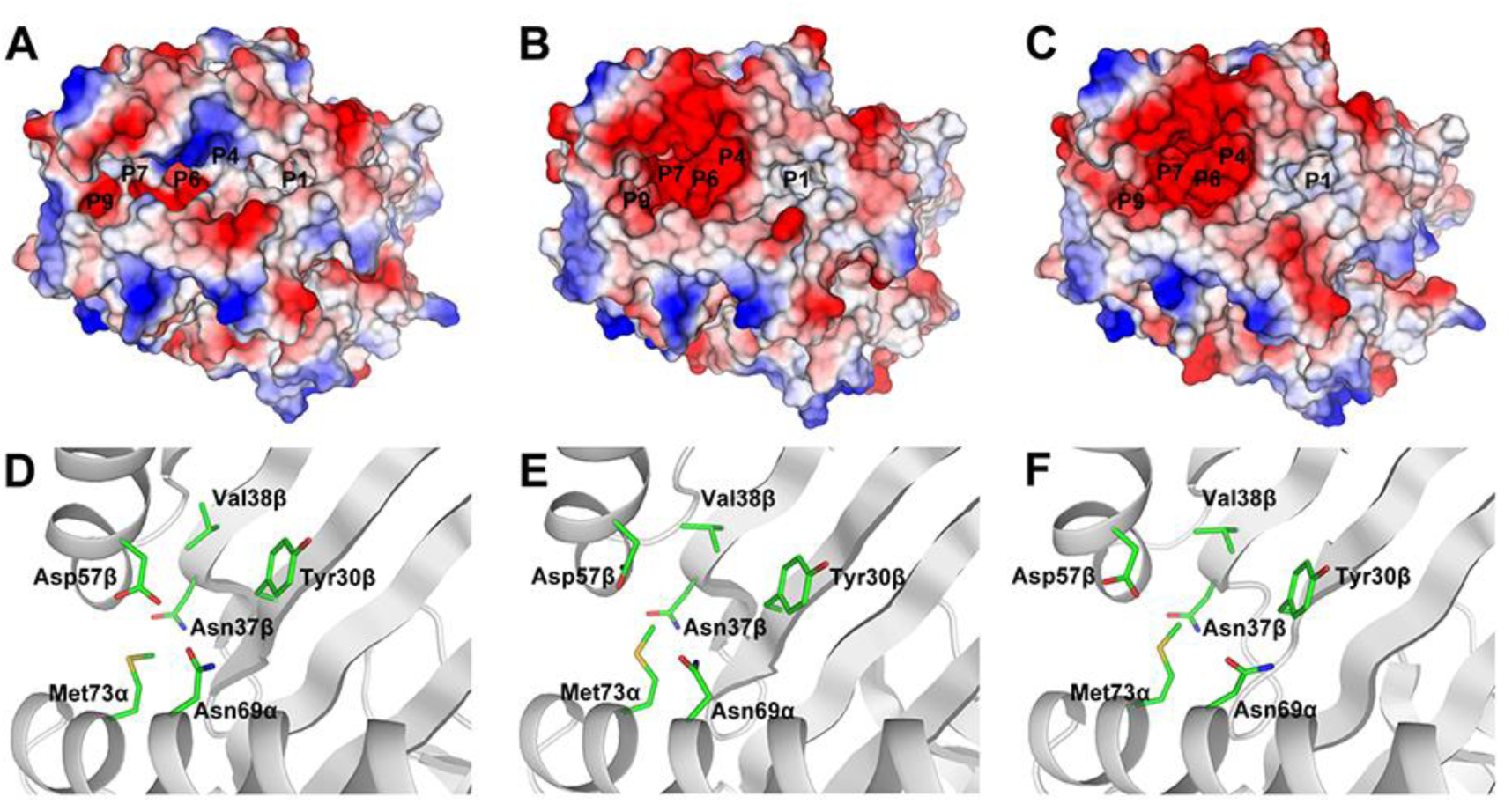
The electrostatic potential surface of *DRA1-DRB1*03:01* (A) *DRA1-DRB1*13:01* (B) and *DRA1-DRB1*13:02* (C). All structure models are displayed by surface. The positive, neutral and negative regions are colored in blue, white and red, respectively. The P9 pocket structure of *DRA1-DRB1*03:01* (D), *DRA1-DRB1*13:01* (E), and *DRA1-DRB1*13:02* (F). HLA molecules and pocket residues are shown as grey cartoon and green sticks, respectively.

## DISCUSSION

In this study, by applying the target capture sequencing, we fine mapped the ACPA-positive RA risks within the MHC region. A key finding of this study is the major influence of HLA-DQα1:160D on ACPA-positive RA instead of the well-described RA-risk *HLA-DRB1* alleles in Han ancestry. HLA-DRβ1:37N is an independent protective factor for ACPA–positive RA. The novel findings are confirmed in an independent case-control cohort by classical HLA genotyping. Furthermore, one of DQα1:160D encoding allele *DQA1*0303* confers high risk for joint damage in patients with smoking in early disease.

Previously, a number of reports have shown that the greatest genetic risk for ACPA-positive RA came from certain *HLA-DRB1* alleles^52-54^, in which the *DRB1*0405*, DRβ1:11, 13, 57, 74, and 71 were reported as strong risks for Asian RA patients^10^. However, even though the classical *HLA-DRB1* alleles have been reported to confer strong RA risks in different populations, other HLA genes have also been shown to associate with ACPA-positive RA. For example, Raychaudhuri *et al*. have reported that single-amino-acid variations in HLA-B (at position 9) and HLA-DPβ1 (at position 9) were strong risks for seropositive RA, besides other three amino acid positions (11, 71 and 74) in HLA-DRβ1^9^. Another study also reported a strong association with ACPA-positive RA at HLA-A amino acid position 77^15^. It has thus been a challenge to draw definitive conclusion concerning the role of different MHC class II alleles in susceptibility to ACPA-positive RA.

In present study, we identified HLA-DQα1:160D as the strongest and independent genetic risk instead of the well-known *HLA-DRB1* alleles for ACPA-positive RA in Han population. Our novel finding could be rationally explained by the fact that DQα1:160D is encoded by two alleles, i.e. *DQA1*0302* and *\*0303*. Though the two alleles are common in East Asians, they are rare variants in Caucasians^55, 56^(http://www.allelefrequencies.net/hla6006a.asp). Thus, the SNPs tagging *DQA1*0302* and *\*0303* may not be included in DNA beadchips for GWAS. This could help to explain why DQα1:160D and its coding alleles *\*0302* and *\*0303* have not been detected or suggested to be risk factors for RA, though many RA GWASs and chip-based genotypic imputations have been performed, including a recent large-scale HLA imputation study of ACPA-positive RA in Japanese population^57^. Furthermore, differences between our findings and previously reported HLA associations could also be due to the progress in HLA typing methodology using sequencing technology that covers the whole MHC class II region, which allows typing of HLA alleles at a much higher resolution than before. Notably, however, in our previous work we have showed that *HLA-DQA1*03* is significantly associated with RA susceptibility (allelic frequencies: 0.351 vs. 0.256, *P* = 4.76 × 10^−4^, OR = 1.56, 95% CI 1.22–2.03)^8^. Consistently with our finding, Raychaudhuri^9^ also reported the allelic frequencies of *DQA1*03* to be increased in seropositive RA patients compared to healthy controls (0.462 vs. 0.185). Moreover, *DQA1*0302* has been reported as a genetic risk for Vitiligo and Ocular myasthenia gravis in Chinese Han population^58, 59^, and for Type 1 diabetes mellitus in children in Japanese population^60^. In agreement with previous studies, our data indicated that *HLA-DRB1*0405*, DRβ1:11, 13, 57, 74, and 71 could be strong risks for ACPA-positive RA, if the *DQA1* association was not noted.

It has been well established that smoking is a risk factor linked to RA susceptibility and severity and this risk is increased by a gene-environment interaction between smoking and *DRB1* alleles and restricted to ACPA-positive RA. Consistent with previous finding^14^, we now also show that *DRB1*0405* carriers with smoking had increased radiographic damage in ACPA-positive RA patients at an early stage of disease. We further show that one of DQα1:160 coding allele *DQA1*0303* has high impact on radiographic severity of ACPA-positive RA, especially in patients with smoking and in early disease. Our data thus supports the notion that smoking, in the presence of RA-risk genetic background, may trigger immunity to citrullinated proteins and lead to RA development and accelerate joint damage.

Although DQα1:160 locate far away from antigen binding groove and may have little influence on epitope binding, it is adjacent to the DQα -DQβ dimer of dimer interface. Previous study has reported that the amino acid substitutions in the dimer of dimer interface of HLA-DRβ1 inhibited CD4^+^ T cell activation^61^. Therefore, we assume that the residue substitutions at DQα1:160 may contribute to the dimer of dimer structure stability and T cell activation. Indeed, by electrostatic interaction analysis we observed a strong electrostatic interaction between negatively charged DQα1 interface residues and positively charged DQβ1’ interface residues. Compared to the non-charged A160α or S160α, the additional negative charge introduced by D160α further enhances the interaction with DQβ1, leading to an increased T cell activation. The DRβ1:37 residue is located within pocket P9 on the beta-sheet floor with their side chains oriented into the peptide-binding groove. Modification of DRβ1:37 residue is sufficient to alter the T-cell receptor peptide recognition^62, 63^. Pocket P9 of class II molecules has been linked to several autoimmune diseases. For example, amino acid variations within the pocket P9 of HLA-DRβ have been shown to associate with increased risk for primary Sjogren’s syndrome^64^. DRβ1:37N was a risk residue for susceptibility to primary sclerosing cholangitis^65^. In present work, by electrostatic potential surface analysis we showed that three DRβ1:37N encoding alleles *\*03:01, *13:01*, and *\*13:02* have negative charged P9 pocket, which benefits electrostatic interaction and could bind with epitope P9 arginine, thus could at least partly explain its protective effect for ACPA-positive RA discovered in present study. In summary, by sequencing of the entire MHC region for discovery and HLA genotyping for validation in two independent cohorts, our study demonstrates that *HLA-DQA1*, instead of *HLA-DRB1*, confers the greatest independent genetic risk for ACPA-positive RA in Chinese Han. Our study also illustrates the value of deep sequencing for fine mapping real risk variants in the MHC region.

## ACKNOWLEDGEMENTS

We thank the staff from Department of Rheumatology and Immunology, People’s Hospital, Peking University, for recruiting patients and healthy controls, the staff from the Computing Platform of the Center for Life Sciences, Peking University, for assisting the functional prediction analysis, and the staff from BGI-Shenzhen who contributed to the technical assistance. We wish to thank the patients and healthy volunteers for their cooperation and for giving consent to participate in this study. This work was supported in part by the National Key Basic Research Program of China (973 Program) (No. 2014CB541901), the National Natural Science Foundation of China (No. 81120108020, No. 31711530023, No. 31670915, No. 31470875, No. 31270914, No. 31530020, No. 31700794, No. 81401329, No. 81771678, No. 81471601, No. 81671604), the National Key Research and Development Program of China (No. 2016YFA05022300), Beijing Natural Science Foundation (No. 7162192), and Shenzhen Municipal of Government of China (No. CXB201108250094A).

## AUTHOR CONTRIBUTIONS

J.G., X.X. and Z.L. conceptualized and designed the study. X.Z and L.K. participated in the study design and supervised manuscript writing. J.G., T.Z. and H.C. coordinated and supervised the study teams. J.G., X.W.L., T.Z., H. L., and H.C. conducted data management and manuscript preparation. T.Z., X.W.L., Y.W.Z., X.M. and H.J.Y. conducted the statistical analyses. H.M.Y., H.J.J., J.W., L.S., L.P., L.H.L. L.L. and K.Y. participated in data interpretation and manuscript writing. M.L., Y.Z., X.S., F.H., Y.D., M.Z., H.J., Xin L., Y.H., Xu L., Y.Y., X.W., X.Z., and Y.S. conducted sample selection and participated in data management. All coauthors edited and reviewed the final version of manuscript.

